# Loop-mediated isothermal amplification assay for *Enterococcus sp., E. coli* and *S. aureus* in chicken

**DOI:** 10.1101/305805

**Authors:** HyeSoon Song, YouChan Bae, HyukMan Kwon, YongKuk Kwon, SeongJoon Joh

## Abstract

Bacterial chondronecrosis with osteomyelitis (BCO) is a major cause of lameness in broiler chicken, and results in serious economic losses worldwide. Although the pathogenesis mechanism leading to lameness is not entirely understood, some strains of *Enterococcus sp*., avian pathogenic *Escherichia coli*, or *Staphylococcus aureus* have been long recognized as important causative pathogens. To prevent the progression of *Enterococcus sp*., avian pathogenic *E. coli*, or *S. aureus* infections, we developed rapid, sensitive, and convenient diagnostic assays using loop-mediated isothermal amplification (LAMP). Entero-Common-LAMP assays were developed for a simultaneous detection of eight *Enterococcus* species. To target specific microorganisms, seven Entero-Specific-LAMP assays for *E. faecalis, E. faecium, E. hirae, E. gallinarum, E. avium, E. durans* and *E. cecorum*, and *E. coli*-LAMP and *S. aureus*-LAMP assays, were developed. Considering the prevalence and economic impact of *Enterococcus sp., E. coli*, and *S. aureus*, the developed ten different LAMP assays have a considerable potential as routine diagnostic methods in the field or in resource-limited environments.

---

Bacterial chondronecrosis with osteomyelitis (BCO) in broiler chicken compromises chicken welfare and causes serious economic losses to the poultry industry worldwide because of reduced chicken productivity and death (12). BCO, including osteomyelitis, femoral head necrosis, long bone necrosis, proximal femoral degeneration, bacterial chondritis with osteomyelitis, and bacterial chondronecrosis, is an important cause of lameness in broiler chicken (13, 18). BCO was first reported in 1972, and the incidence of lameness with BCO has increased significantly in Australia, USA, Canada, and Europe over the past two decades, with recent reports indicating that over 1% of all broilers grown to heavy processing weights may be affected after 5 wk of age (12, 25, 29). An investigation in Bulgaria revealed the significant scale of the problem, with lameness accounting for 10% of mortality in lame chickens, and BCO accounting for more than 90% of these cases (5). Although the complex pathogenicity mechanism of BCO is not entirely understood, *Enterococcus sp*., avian pathogenic *Escherichia coli*, and *Staphylococcus aureus* are recognized important pathogens associated with BCO (5, 7, 17, 23, 29–30). These bacteria are ubiquitous in poultry environments where the birds are hatched, reared, or processed; they are transmitted to chicks from breeder parents, contaminated eggs, or hatchery sources by opportunistic infection (18, 29). Further, BCO appears to occur when *Enterococcus sp., E. coli*, or *S. aureus* infects the broilers via the integument, respiratory system, or gastrointestinal tract; circulates in the bloodstream; and forms micro-abscesses that cause infarction and local metaphyseal bone necrosis (17, 29). The condition of broiler chicken affected by BCO usually progresses fairly rapidly from mild to severe lameness.

Development of a rapid and specific method for the detection of *Enterococcus sp., E. coli*, or *S. aureus* infection in the field and in resource-limited environments is important for the prevention of the progression of BCO. Loop-mediated isothermal amplification (LAMP) is a highly specific, efficient, and rapid method based on 2-3 sets of primers that target a gene under isothermal conditions, with no special equipment for DNA amplification required. After the LAMP reaction, a positive result is detected by assessing increase in sample turbidity (determined using a real-time turbidity meter) caused by the formation of a magnesium pyrophosphate byproduct, or by visual inspection (color change); there is no need for agarose gel electrophoresis (6, 19, 21).

In the current study, by targeting highly conserved genes of BCO-associated bacteria, we successfully developed an Entero-Common-LAMP assay for a simultaneous detection of common enterococcus gene of eight *Enterococcus sp. (E. faecalis, E. faecium, E. hirae, E. gallinarum, E. avium, E. durans, E. cecorum*, and *E. columbae)*, seven types of Entero-Specific-LAMP assays for a specific detection of seven *Enterococcus sp. (E. faecalis, E. faecium, E. hirae, E. gallinarum, E. avium, E. durans*, and *E. cecorum)*; and *E.coli*-LAMP and *S. aureus*-LAMP assays.

## MATERIALS AND METHODS

### Bacterial and viral strains

*Enterococcus* strains *E. faecalis* (ATCC 29212), *E. faecium* (ATCC 19434), *E. hirae* (ATCC 8043), *E. gallinarum* (ATCC 49573), *E. avium* (ATCC 14025), *E. durans* (ATCC 19432), *E. cecorum* (ATCC 43198), *E. columbae* (ATCC 51263), *E. mundtii* (ATCC 43186), *E. saccharolyticus* (ATCC 43076), *E. casseliflavus* (ATCC 25788), and *E. sulfureus* (ATCC 49903); *Escherichia coli* (ATCC 25922); *Staphylococcus* strains *S. aureus* (ATCC 25923), *S. cohnii* (ATCC 35662), *S. xylosus* (ATCC 29971), *S. lentus* (ATCC 49574), *S. hominis* (field isolate), and *S. epidermidis* (field isolate); *Ornithobacterium rhinotracheale* (field isolate); *Pasteurella multocida* (field isolate); *Mycoplasma gallisepticum* (ATCC 19610); *Mycoplasma synoviae* (ATCC 25204); *Bacillus cereus* (ATCC 14579); *Campylobacter coli* (ATCC 33559); *Clostridium perfringens* (ATCC 13124); *Campylobacter jejuni* (ATCC 33560); *Salmonella enteritidis* (ATCC 31194); chicken infectious anemia virus (CIAV, field isolate); reticuloendotheliosis virus (REV, field isolate); and Marek’s disease virus (MDV, ATCC VR-624™) were from the American Type Culture Collection, and used as reference strains in the current study.

### Isolation of DNA

Total bacterial and viral DNA was extracted from the microorganisms using QIAamp DNA mini kit (Qiagen, Germany) and QIAamp DNeasy Blood and Tissue kit (Qiagen), according to the manufacturer’s instructions.

### Design of universal and species-specific LAMP primers

Entero-Common-LAMP primers for a simultaneous detection of *E. faecalis, E. faecium, E. hirae, E. gallinarum, E. avium, E. durans, E. cecorum*, and *E. columbae* were designed based on the published *rpoB* gene sequences of the *Enterococcus sp*. Additional LAMP primers, specific to *E. faecalis, E. faecium, E. hirae, E. gallinarum, E. avium, E. cecorum*, and *E. durans*, were designed to target specific variable regions by using Primer Explorer V4 software (Eiken Chemical Co., Ltd., Japan). The *malB* and *nuc* genes were selected as target genes for the detection of *E. coli* and *S. aureus*. These LAMP primers included the following: a forward outer primer F3; a reverse outer primer B3; a forward inner primer FIP (harboring the F2 region sequence at its 3’-end and the F1c region sequence at its 5’-end); a reverse inner primer BIP (harboring the B2 region sequence at its 3’-end and the B1c region sequence at its 5’-end); a forward loop primer LF; and a reverse loop primer LB. These primers recognized eight conserved regions within their target genes (Table 1).

**TABLE 1.**
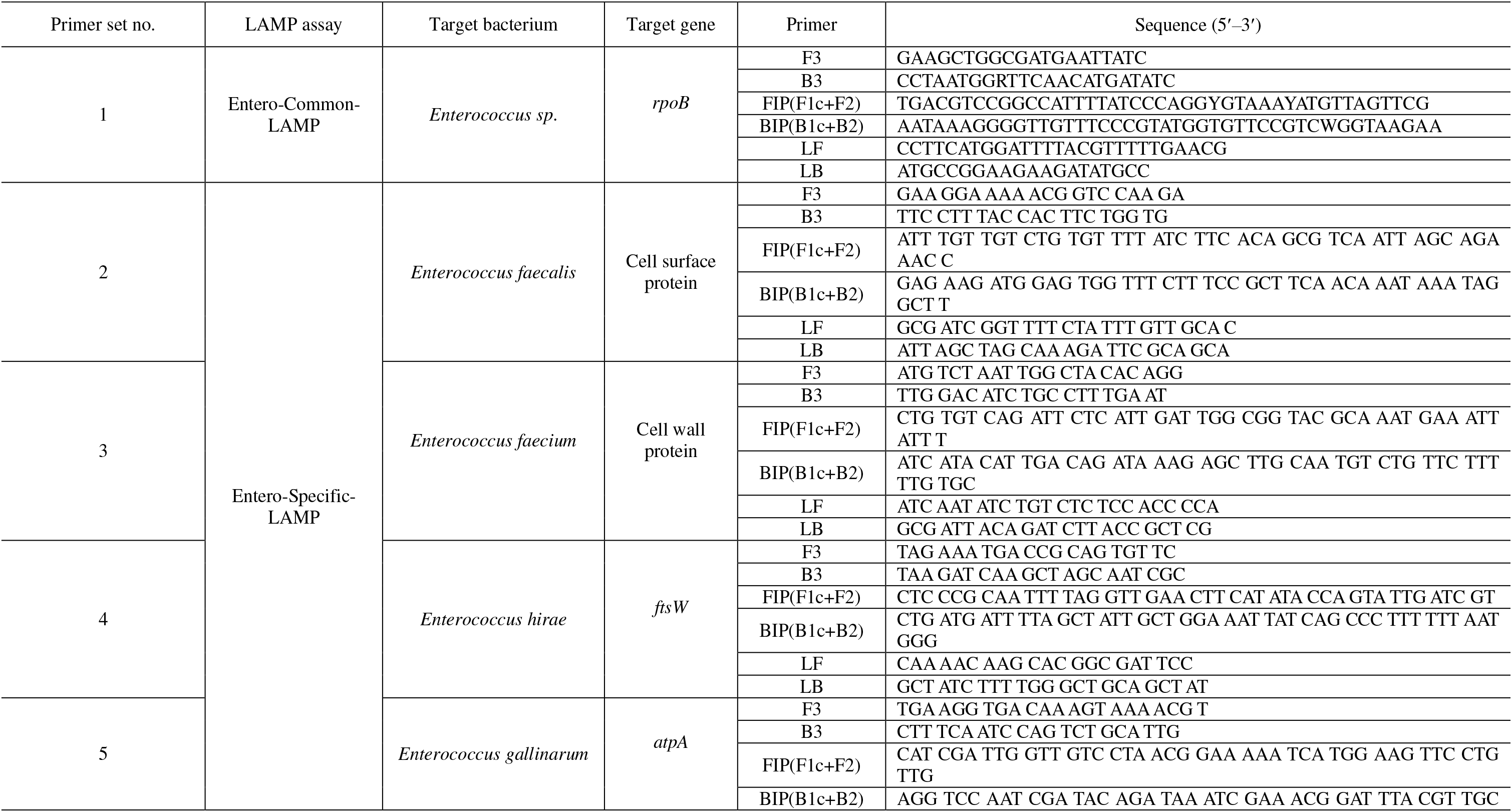
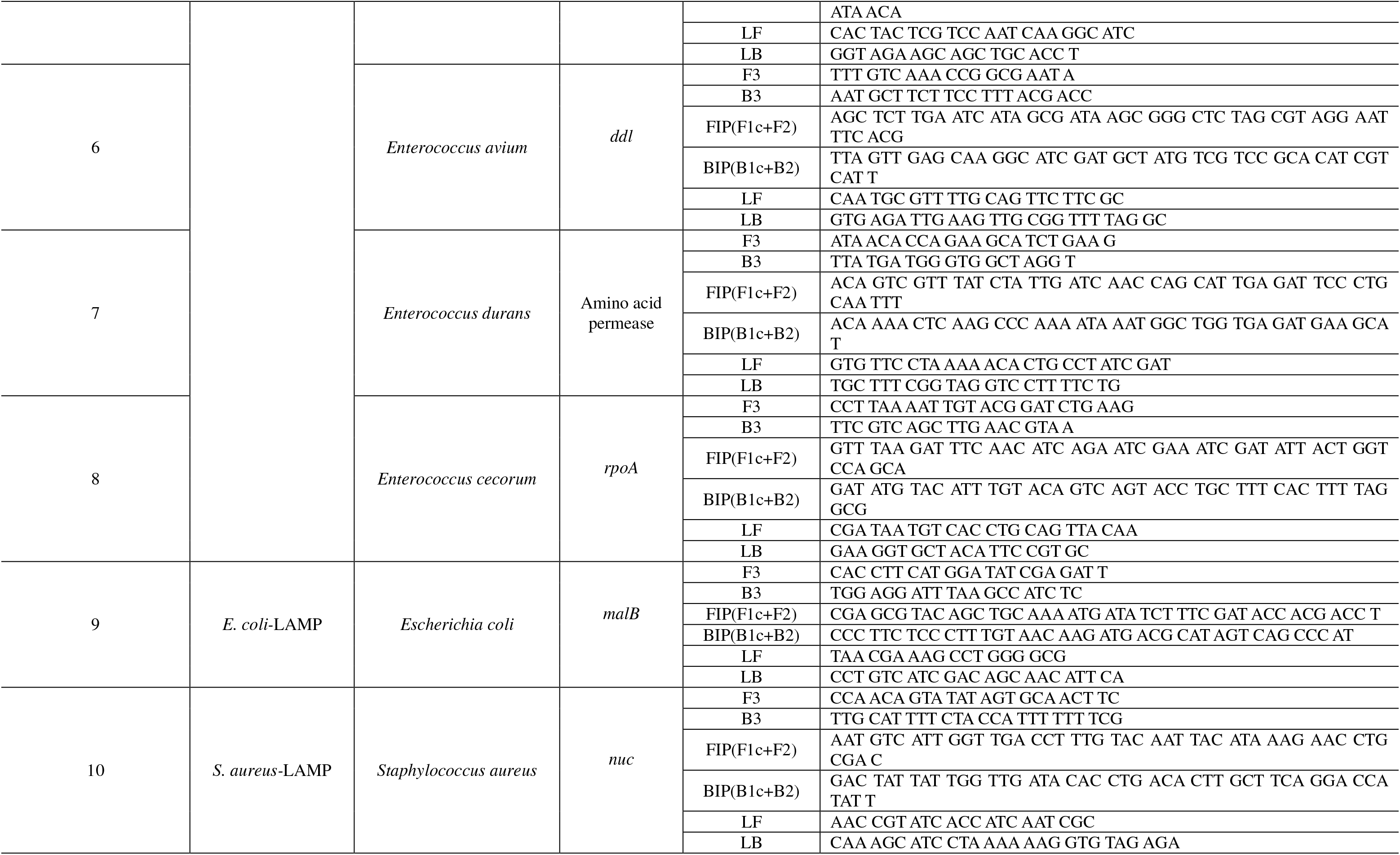
LAMP primers for the detection of *Enterococcus sp*., *E. coli*, and *S. aureus*

### LAMP assays

The LAMP reactions were performed in 25 μl reaction volumes using a Mmiso DNA amplification kit (Mmonitor, South Korea), in accordance with the manufacturer’s instructions. The reaction mixtures contained 1 μl of bacterial genomic DNA, 2× reaction buffer, 8 U of *Bst* DNA polymerase, 2.5 pmol of the outer primers (F3 and B3), 20 pmol of the forward and reverse inner primers (FIP and BIP), and 10 pmol of the loop primers (LF and LB). The LAMP assays were performed under isothermal conditions at 63°C for 30 min, followed by heating to 80°C for 5 min in a heating block, to terminate the reaction.

### The specificity and detection limits of LAMP

The specificities of the optimized Entero-Common-LAMP, seven types of Entero-Specific-LAMP assays (i.e., *E. faecalis-LAMP, E. faecium-LAMP, E. hirae-LAMP, E. gallinarum-LAMP, E. avium*-LAMP, *E. durans*-LAMP, and *E. cecorum*-LAMP), *E. coli*-LAMP and *S. aureus*-LAMP assays were tested using all bacteria and viruses. Each LAMP reaction was performed using 25 ng of genomic DNA, at 63°C for 30 min, and terminated by heating to 80°C for 5 min in a heating block. The assay detection limits were determined by testing 5-fold serial dilutions of bacterial DNA of *E. faecalis, E. faecium, E. hirae, E. gallinarum, E. avium, E. durans, E. cecorum, E. columbae, E. coli*, and *S. aureus*.

### PCR

To compare the detection limits of the PCR and LAMP assays, the target genes from *E. faecalis, E. faecium, E. hirae, E. gallinarum, E. avium, E. durans, E. cecorum, E. columbae, E. coli*, and *S. aureus* were PCR-amplified in individual reactions. The reaction volume was 20 μl, and the reactions contained 2.5 mM each dNTP, 1.5 mM MgCl_2_, 10× reaction buffer, 1 U of Taq polymerase, 10 pM LAMP F3 and B3 primers, and 1 μl of serial dilutions of template DNA. The thermal cycler (Eppendorf, Germany) was set to the following PCR conditions: 94°C for 5 min; 30 cycles of 94°C for 45 s, 55°C for 45 s, and 72°C for 45 s; with a final elongation at 72°C for 7 min. For the Entero-common-PCR with *E. gallinarum* and *E. avium*-specific-PCR of *E. avium*, the reactions conditions were as follows: an initial denaturation at 94°C for 5 min; 40 cycles of denaturation at 94°C for 1 min, annealing at 53°C for 1 min, and polymerization at 72°C for 1 min; and an extension at 72°C for 7 min. The PCR products were analyzed under UV light after 1.5% agarose gel electrophoresis.

### Detection of *Enterococcus sp*., *E. coli*, and *S. aureus* in field samples

The Entero-Common-LAMP, seven types of Entero-Specific-LAMP, *E. coli*-LAMP, and *S. aureus*-LAMP assays were used to analyze *E. faecalis, E. faecium, E. hirae, E. gallinarum, E. avium, E. cecorum, E. coli*, and *S. aureus* isolated from the liver, femur, and joint of broiler chickens with lameness. The isolated *Enterococcus sp., E. coli*, and *S. aureus* strains were cultivated on tryptic soy agar. Genomic DNA was extracted and used in the nine types of LAMP assays to compare the results of *16S rRNA* sequencing, PCR, and LAMP assays.

## RESULTS AND DISCUSSION

Several LAMP primer sets targeting specific genes (including *rpoB, ftsW, atpA, rpoA*, and *ddl* of *Enterococcus sp.; malB* of *E. coli*; and *nuc* of *S. aureus)* were screened using a DNA amplification kit according to the manufacturer’s instructions. After the LAMP amplification under isothermal conditions, a color change from violet to sky blue indicated a positive reaction and the negative reaction remained violet. The sequences of the optimal LAMP primer sets are shown in Table 1. The specificity of optimal LAMP primer sets was examined using 25 ng of genomic DNA extracted from 28 different bacteria representing various genera and species, and three viruses. As shown in Fig. 1A, only in tubes containing 8 strains of *Enterococcus* genomic DNA and specific primers, the reaction mixture color changed from violet to sky blue, while the mixtures in other tubes remained violet. Likewise, the LAMP primers specific to *E. coli, S. aureus, E. faecalis, E. faecium, E. hirae, E. gallinarum, E. avium, E. durans*, and *E. cecorum* yielded amplification products only in the reaction tubes that contained the specific target genomic DNA (Fig. 1C and E; Fig. 2A, C, E, G, I, K, and M). The positive reactions were also confirmed by the presence of ladder-like DNA bands on 1.5% TAE agarose gels (Fig. 1D and F; Fig. 2B, D, F, H, J, L, and N).

**FIG 1.**
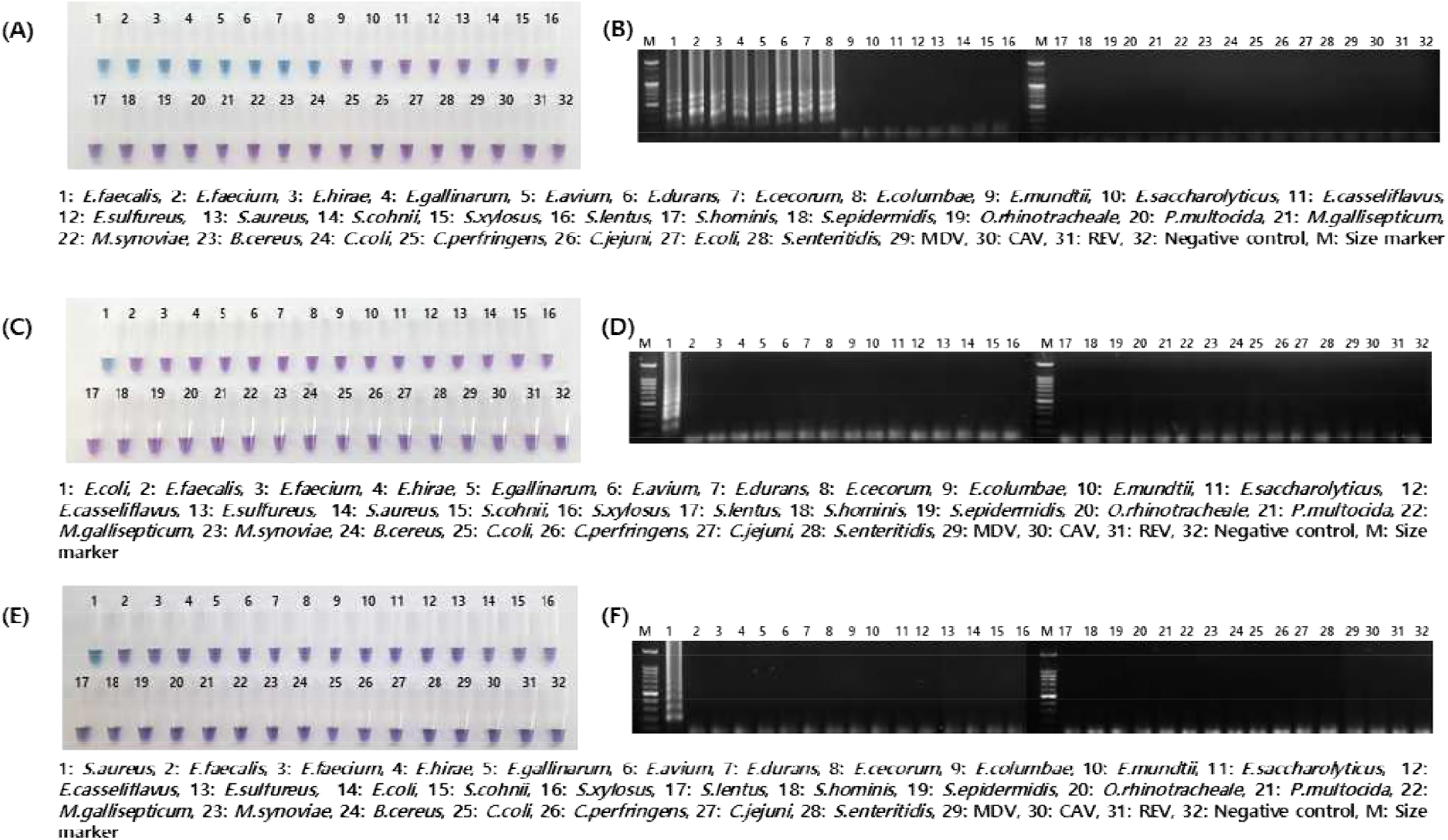
Specificity of the Entero-Common-LAMP, *E. coli*-LAMP, and *S. aureus*-LAMP assays. (A) Entero-Common-LAMP; (C) *E. coli*-LAMP; and (E) *S. aureus*-LAMP. A color change from violet to sky blue indicated a LAMP-positive reaction, while the color of a LAMP-negative reaction mixture remained violet. The LAMP products were also resolved by 1.5% agarose gel electrophoresis: (B) Entero-Common-LAMP; (D) *E. coli*-LAMP; and (F) *S. aureus*-LAMP.

**FIG 2.**
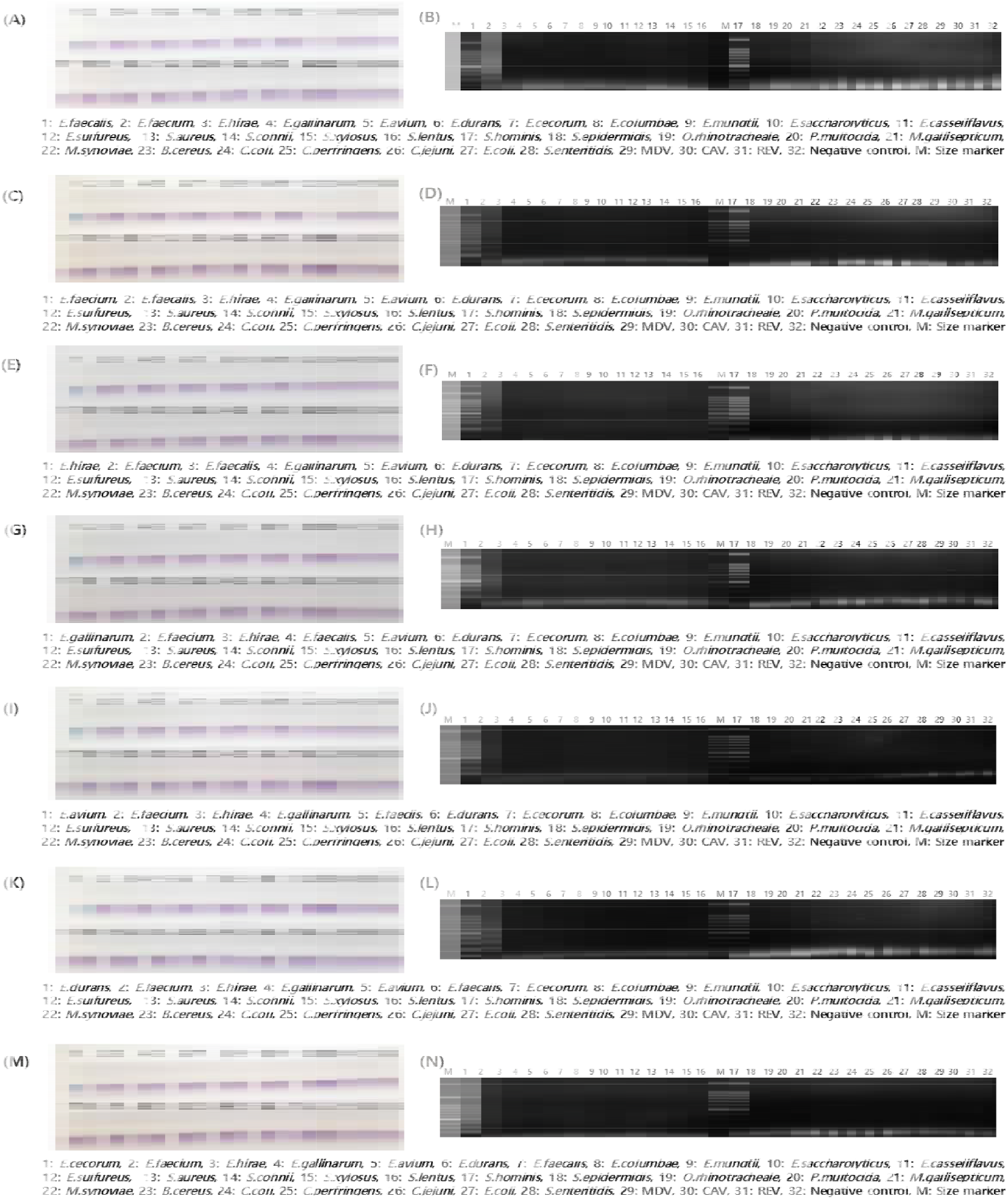
Specificity of seven types of the Entero-Specific-LAMP assays. (A) *E. faecalis*-LAMP; (C) *E. faecium*-LAMP; (E) *E. hirae*-LAMP; (G) *E. gallinarum*-LAMP; (I) *E. avium*-LAMP; (K) *E*. durans-LAMP; and (M) *E. cecorum*-LAMP. In these assays, a color change from violet to sky blue was observed only in the tubes containing the target genomic DNA. The products of the LAMP assays were resolved by 1.5% agarose gel electrophoresis: (B) *E. faecalis*-LAMP; (D) *E. faecium*-LAMP; (F) *E*. hirae-LAMP; (H) *E. gallinarum*-LAMP; (J) *E. avium*-LAMP; (L) *E*. durans-LAMP; and (N) *E. cecorum*-LAMP.

To compare the detection limits of the ten types of LAMP assays and that of conventional PCR, 5-fold serially diluted genomic DNA samples from *E. faecalis, E. faecium, E. hirae, E. gallinarum, E. avium, E. durans, E. cecorum, E. columbae, E. coli*, and *S. aureus* were used. The evaluation of the reaction detection limits was performed using primers LAMP F3 and B3 by PCR(Table 1). As shown in Fig. 3, the detection limits for the Entero-Common-LAMP assay were 2 pg/μl for *E. faecalis* and *E. durans;* 10 pg/μl for *E. faecium* and *E. columbae;* 400 fg/μl for *E. hirae;* 40 pg/μl for *E. gallinarum;* 4 pg/μl for *E. avium;* and 50 pg/μl for *E. cecorum*. Therein, the success of the LAMP reaction was detected with the naked eye and by agarose gel electrophoresis. On the other hand, the detection limits of PCR using the Entero-Common-LAMP primers F3 and B3 were 50 pg/μl for *E. faecalis, E. faecium, E. durans*, and *E. columbae;* 1.25 ng/μl for *E. hirae;* 250 pg/μl for *E. gallinarum;* 10 pg/μl for *E. avium;* and 250 pg/μl for *E. cecorum* (Fig. 5A-H).

**FIG 3.**
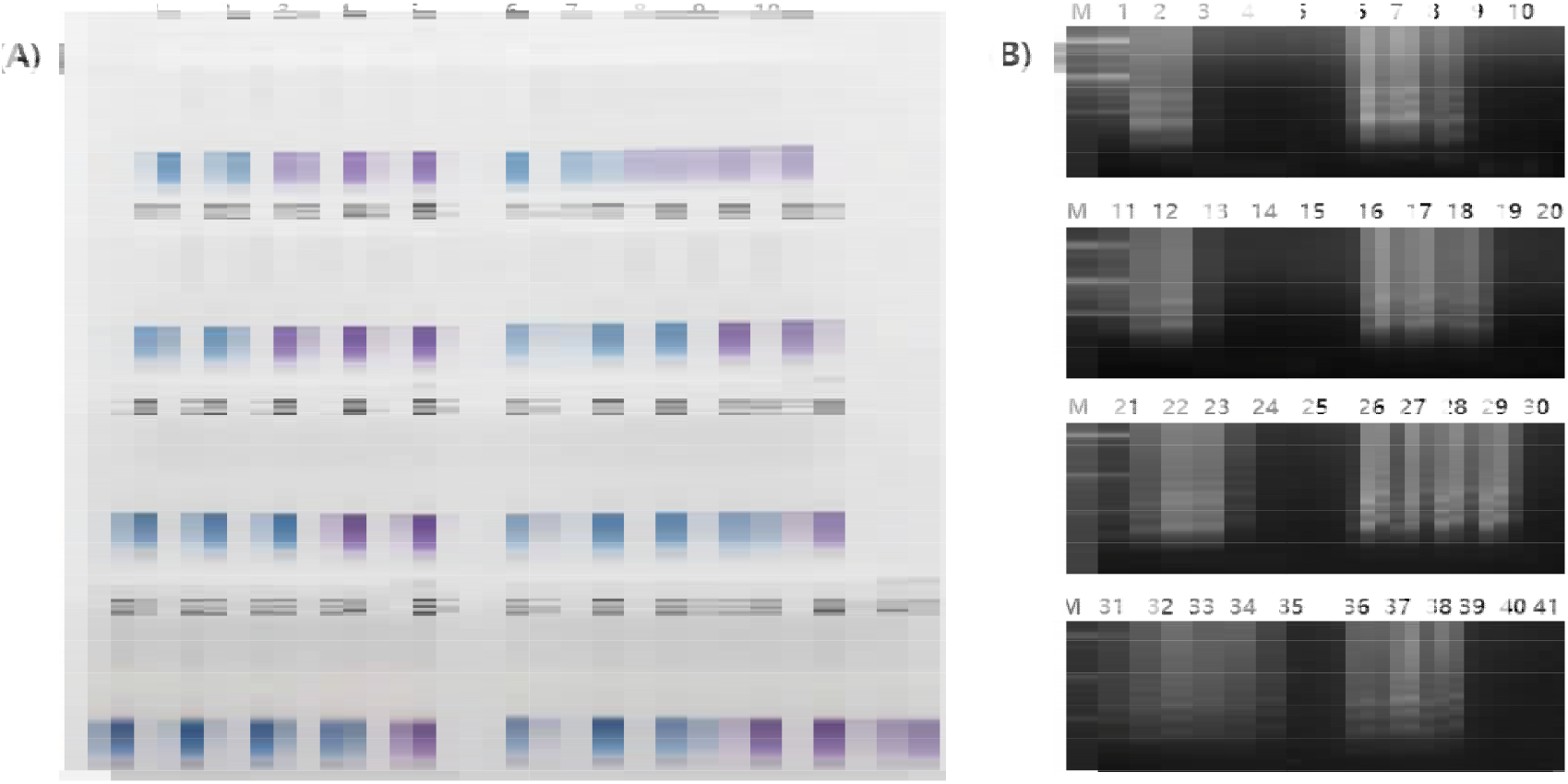
Detection limits of the Entero-Common-LAMP assay. (A) Naked-eye visualization of the LAMP products. The color of LAMP-positive reactions turned sky blue, while the color of LAMP-negative reactions remained violet. (B) Agarose gel electrophoresis of LAMP products. Lane M, 100 bp DNA marker. *E. faecalis* genomic DNA (lanes/tubes): 1, 10 pg/μl; 2, 2 pg/μl; 3, 400 fg/μl; 4, 80 fg/μl; and 5, 16 fg/μl. *E. faecium* genomic DNA (lanes/tubes): 6, 50 pg/μl; 7, 10 pg/μl; 8, 2 pg/μl; 9, 400 fg/μl; and 10, 80 fg/μl. *E. durans* genomic DNA (lanes/tubes): 11, 10 pg/μl; 12, 2 pg/μl; 13, 400 fg/μl; 14, 80 fg/μl; and 15, 16 fg/μl. *E. hirae* genomic DNA (lanes/tubes): 16, 10 pg/μl; 17, 2 pg/μl; 18, 400 fg/μl; 19, 80 fg/μl; and 20, 16 fg/μl. *E. columbae* genomic DNA (lanes/tubes): 21, 250 pg/μl; 22, 50 pg/μl; 23, 10 pg/μl; 24, 2 pg/μl; and 25, 400 fg/μl. *E. avium* genomic DNA (lanes/tubes): 26, 500 pg/μl; 27, 100 pg/μl; 28, 20 pg/μl; 29, 4 pg/μl; and 30, 800 fg/μl. *E. gallinarum* genomic DNA (lanes/tubes): 31, 5 ng/μl; 32, 1 ng/μl; 33, 200 pg/μl; 34, 40 pg/μl; and 35, 8 pg/μl. *E. cecorum* genomic DNA (lanes/tubes): 36, 1.25 ng/μl; 37, 250 pg/μl; 38, 50 pg/μl; 39, 10 pg/μl; and 40, 2 pg/μl. Lane 41, negative control.

Seven types of Entero-Specific-LAMP assays detected the target genes from 400 fg/μl *E. faecalis, E. faecium, E. hirae*, and *E. avium* DNA, and 2 pg/μl *E. gallinarum, E. durans*, and *E. cecorum* DNA, under isothermal conditions, within 30 min, and assessment by the naked eye (Fig. 4). The detection limits of PCR with the Entero-Specific-LAMP primers F3 and B3 were 250 pg/μl for *E. faecalis, E. faecium, E. gallinarum*, and *E. durans;* 50 pg/μl for *E. hirae* and *E. cecorum*; and 6.25 ng/μl for *E. avium* (Fig. 5I-O). The detection limits for *E. coli* and *S. aureus* using LAMP assays were 2 pg/μl and 400 fg/μl DNA, respectively; however, the detection limits of PCR were 50 pg/μl for *E. coli* and 10 pg/μl for *S. aureus* (Fig. 5P and Q).

**FIG 4.**
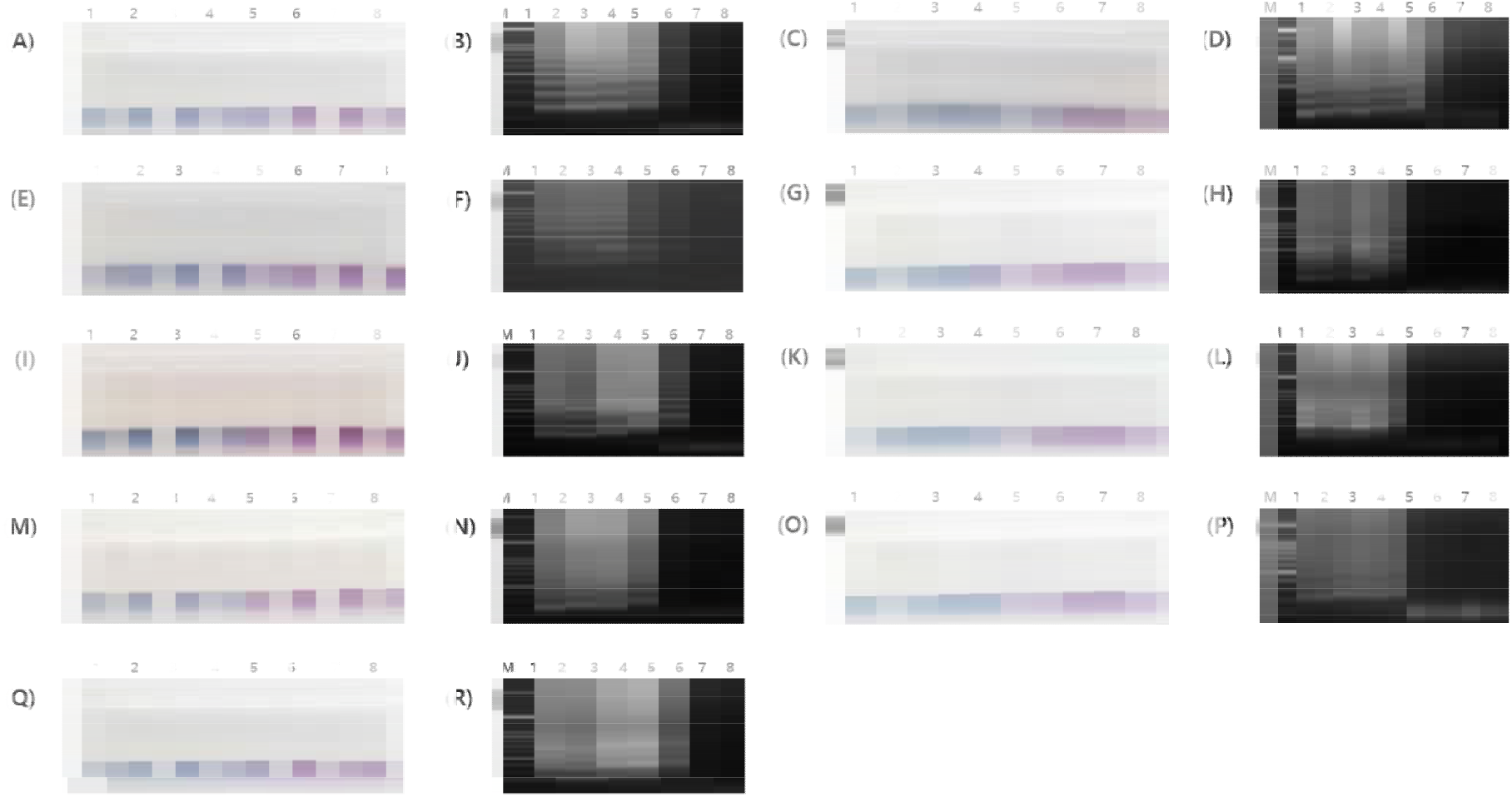
Detection limits of seven types of the Entero-Specific-LAMP, *E. coli*-LAMP, and *S. aureus*-LAMP assays. Visual inspection of LAMP products for the detection of *E. faecalis* (A), *E. faecium* (C), *E. hirae* (E), *E. galliinarum* (G), *E. avium* (I), *E. durans* (K), *E. cecorum* (M), *E. coli* (O), and *S. aureus* (Q) under natural light. Agarose gel electrophoresis of LAMP products from different LAMP assays: *E. faecalis*-LAMP (B), *E. faecium*-LAMP (D), *E. hirae*-LAMP (F), *E. galliinarum*-LAMP (H), *E. avium*-LAMP (J), *E. durans*-LAMP (L), *E. cecorum*-LAMP (N), *E.* coli-LAMP (P), and *S. aureus*-LAMP (R). Lane M, 100 bp DNA marker; lanes (tubes): 1, 250 pg/μl; 2, 50 pg/μl; 3, 10 pg/μl; 4, 2 pg/μl; 5, 400 fg/μl; 6, 80 fg/μl; 7, 16 fg/μl; and 8, negative control.

**FIG 5.**
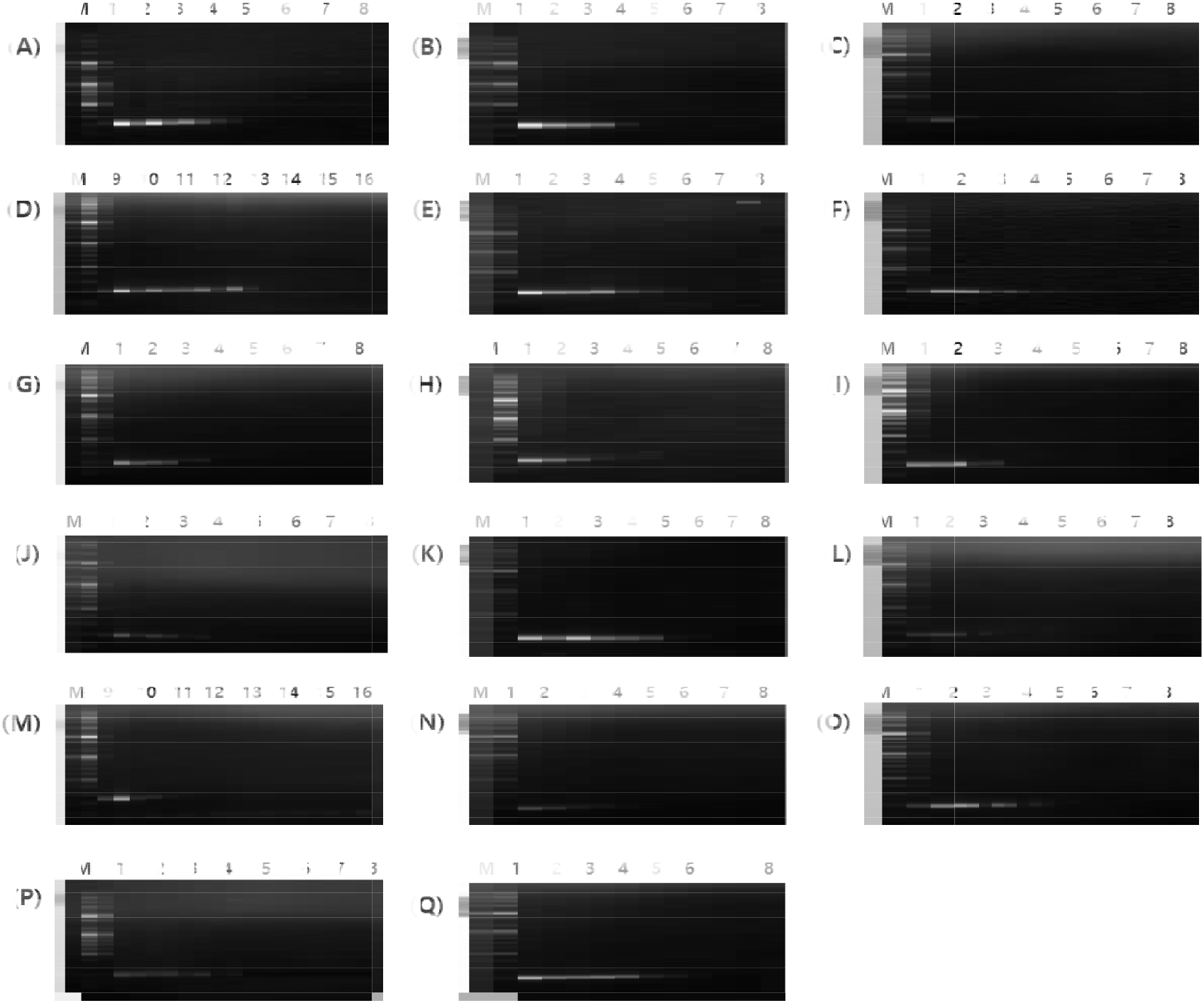
Electrophoretic analysis of PCR products to compare the detection limits of conventional PCR and LAMP assays. PCR was performed to detect *E. faecalis* (A), *E. faecium* (B), *E. hirae* (C), *E. gallinarum* (D), *E. avium* (E), *E. durans* (F), *E. cecorum* (G), and *E. columbae* (H), using universal primers F3 and B3 for *Enterococcus* species. Gels in (I-Q) show resolved PCR products of specific target genes from *E. faecalis* (I), *E. faecium* (J), *E. hirae* (K), *E. gallinarum* (L), *E. avium* (M), *E. durans* (N), *E. cecorum* (O), *E. coli* (P), and *S. aureus* (Q). Lanes: M, 100 bp DNA marker; 1, 6.25 ng/μl; 2, 1.25 ng/μl; 3, 250 pg/μl; 4, 50 pg/μl; 5, 10 pg/μl; 6, 2 pg/μl; 7, 400 fg/μl; 9, 31.3 ng/μl; 10, 6.25 ng/μl; 11, 1.25 ng/μl; 12, 250 pg/μl; 13, 50 pg/μl; 14, 10 pg/μl; 15, 2 pg/μl; 8 and 16, negative control.

In conclusion, the sensitivity of the Entero-Common-LAMP and Entero-Specific-LAMP assays (for *E. faecalis, E. faecium, E. hirae, E. gallinarum, E. avium, E. durans*, and *E. cecorum*), and *E. coli*-LAMP and *S. aureus*-LAMP assays, was higher than that of conventional PCR for detecting pathogens associated with the lameness in broiler chicken.

In total, 140 bacterial strains isolated between 2016 and 2017 from the liver, femur, and joint of broiler chickens with lameness from the Animal and Plant Quarantine Agency in Korea were analyzed by LAMP assays, conventional PCR, and *16S rRNA* sequencing. The nine types of LAMP assays and PCR were congruent (100%). The agreement between the LAMP assays and *16S rRNA* sequencing was 92.6%, 83.9%, 95.2%, 0%, 100%, and 95% for the detection of *E. faecalis, E. faecium, E. hirae, E. gallinarum, E. avium*, and the avian pathogenic *E. coli*, respectively (Table 2).

**TABLE 2.** Outcomes of nine types of LAMP assays of clinical samples, compared with the diagnostic PCR and *16S rRNA* sequencing assays^a^

| Species | LAMP |  |  |  | PCR |  |  |  | *16S rRNA* sequencing |  |  |  |
| --- | --- | --- | --- | --- | --- | --- | --- | --- | --- | --- | --- | --- |
|  | P/T | N/T | F/T | Sensitivity | P/T | N/T | F/T | Sensitivity | P/T | N/T | F/T | Sensitivity |
| *Enterococcus sp. common* | 87/140 | 53/140 | 0/140 | 100% | 87/140 | 53/140 | 0/140 | 100% | NA | NA | NA | NA |
| *E. faecalis* | 27/140 | 113/140 | 0/140 | 100% | 27/140 | 113/140 | 0/140 | 100% | 25/140 | 113/142 | 2/140 | 92.6% |
| *E. faecium* | 27/140 | 113/140 | 0/140 | 100% | 27/140 | 113/140 | 0/140 | 100% | 26/140 | 109/140 | 5/140 | 83.9% |
| *E. hirae* | 20/140 | 120/140 | 0/140 | 100% | 20/140 | 120/140 | 0/140 | 100% | 20/140 | 119/140 | 1/140 | 95.2% |
| *E. gallinarum* | 9/140 | 131/140 | 0/140 | 100% | 9/140 | 131/140 | 0/140 | 100% | 0/140 | 131/140 | 9/140 | 0% |
| *E. avium* | 1/140 | 139/140 | 0/140 | 100% | 1/140 | 139/140 | 0/140 | 100% | 1/140 | 139/140 | 0/140 | 100% |
| *E. cecorum* | 3/140 | 137/140 | 0/140 | 100% | 3/140 | 137/140 | 0/140 | 100% | NA | NA | NA | NA |
| *E. coli* | 38/140 | 102/140 | 0/140 | 100% | 38/140 | 102/140 | 0/140 | 100% | 38/140 | 100/140 | 2/140 | 95% |
| *S. aureus* | 10/140 | 130/140 | 0/140 | 100% | 10/140 | 130/140 | 0/140 | 100% | NA | NA | NA | NA |

<sup>a</sup>P, number of true positives; T, number of total samples; N, number of true negatives; F, number of false positives and false negatives; NA, not applicable.

Lameness with BCO is an important skeletal disease of broiler chicken (20). Some strains of *Enterococcus sp*., avian pathogenic *E. coli*, and *S. aureus* are recognized as important BCO pathogens in the poultry industry, and are isolated from single or mixed cultures from BCO lesions (1, 28). In addition, *E. cecorum* and *E. hirae* are recovered from the joint and femur of lame birds (8, 11, 13, 25, 31). Enterococci, including *E. faecalis, E. faecium, E. avium, E. durans*, and *E. gallinarum*, are frequently isolated from the litter, feed, dead-shell, or 1-day-old chicks in poultry farms (2, 4). In one study, avian pathogenic *E. coli* was recovered from over 90% of bacteriologically tested chickens with lameness (5), and was the most frequently isolated bacterium from chickens with BCO (30). *S. aureus* is the major pathogen responsible for bone and joint infections (16), and is also isolated from the litter, feeders, drinkers, and the air in poultry houses (17, 24, 26). *Enterococcus sp*., avian pathogenic *E. coli*, and *S. aureus* are also responsible for significant financial losses worldwide. Therefore, they should be detected precisely and as early as possible to eradicate them and prevent their transmission. To achieve this, a simple and rapid diagnostic method for the detection of *Enterococcus sp., E. coli*, and *S. aureus* in broiler chicken with lameness is necessary. At present, MALDI-TOF, VITEK, and *16S rRNA* sequencing analyses following bacterial isolation are routinely used for the identification of *Enterococcus sp., E. coli*, and *S. aureus*. Although bacterial identification after isolation is the most reliable method, it is time-consuming and labor intensive. PCR-based methods are well optimized with respect to the sensitivity, specificity, and repeatability of the amplification of a target gene, and detect pathogens more quickly than bacterial culture. However, these methods require special equipment, such as thermal cyclers and skilled labor, and PCR amplicons must be analyzed by electrophoresis (22). By contrast, LAMP is a simple, rapid, efficient, and cost-effective method, which uses a water bath or block heater to amplify the target DNA under isothermal conditions. The success of the LAMP amplification reaction may be assessed by the naked eye, either as a turbidity change (white precipitate formation), or through a color change (from violet to sky blue), without the need for electrophoretic analysis.

In the current study, we developed different types of LAMP assays to detect *Enterococcus sp., E. coli*, and *S. aureus*. We designed ten sets of primers (six primers each) targeting the *rpoB* gene from eight common *Enterococcus* species; seven *Enterococcus-specific* genes, i.e., a cell surface protein gene of *E. faecalis*, a cell wall protein gene of *E. faecium*, the *ftsW* gene of *E. hirae*, the *atpA* gene of *E. gallinarum*, the *ddl* gene of *E. avium*, an amino acid permease gene of *E. durans*, and the *rpoA* of *E. cecorum*; the *malB* gene of *E. coli*; and the *nuc* gene of *S. aureus*. We then tested the reaction detection limits and specificity in LAMP reactions performed at 63°C for 30 min.

The detection limit of the Entero-Common-LAMP assay was between 50 pg/μl and 400 fg/μl, whereas the detection limit of the conventional PCR using the Entero-Common-LAMP primers F3 and B3 was between 1.25 ng/μl and 10 pg/μl. This demonstrated that the Entero-Common-LAMP assay was 5-10 times more sensitive than the Entero-common-PCR. Specifically, in the case of *E. hirae*, the detection limits of Entero-Common-LAMP assay and Entero-common-PCR were 400 fg/μl and 1.25 ng/μl, respectively, which indicated that LAMP was 3,125 times more sensitive than the PCR reaction. Further, seven types of Entero-Specific-LAMP assays detected the target genes from 400 fg/μl *E. faecalis, E. faecium, E. hirae*, and *E. avium* DNA; and from 2 pg/μl *E. gallinarum, E. durans*, and *E. cecorum* DNA. By contrast, the detection limits of PCR with the Entero-Specific-LAMP primers F3 and B3 were 250 pg/μl, for *E. faecalis, E. faecium, E. gallinarum*, and *E. durans*; 50 pg/μl, for *E. hirae* and *E. cecorum*; and 6.25 ng/μl, for *E. avium*. The sensitivity of the Entero-Specific-LAMP assays was therefore 25-625 times higher than that of the Entero-specific-PCR reactions. Above all, the sensitivity of the LAMP assay for the detection of *E. avium* was 15,625 times higher than that of the PCR reaction. The detection limits of *E. coli*-LAMP and *S. aureus*-LAMP were 2 pg/μl and 400 fg/μl, respectively; however, the detection limits of the PCR reactions were 50 pg/μl for *E. coli* and 10 pg/μl for *S. aureus*. Furthermore, the sensitivities of the *E. faecalis*-LAMP and *S. aureus*-LAMP assays were superior to those reported previously (9, 14–15, 27, 32). Collectively, these observations indicated that the sensitivity of the ten LAMP assays was much higher than that of conventional PCR and of previously devised LAMP assays.

The specificity tests for the Entero-Common-LAMP, the seven types of Entero-Specific-LAMP, and the *E. coli*-LAMP and *S. aureus*-LAMP assays revealed that the target genes were successfully detected without cross-reactivity with other avian bacterial and viral pathogens.

The practical application of the LAMP assays was evaluated using 140 samples, and the outcomes were compared with those of PCR and *16S rRNA* sequencing. The seven LAMP assays and PCR reactions accurately identified all samples of different *Enterococcus* isolates (including *E. faecalis, E. faecium, E. hirae, E. avium*, and *E. cecorum)* at the genus and species level. Further, the LAMP and PCR assays were 100% congruent for both *S. aureus* and *E. coli* detection. By contrast, the results of *16S rRNA* sequencing indicated 92.6%, 83.9%, 95.2%, and 100% agreement in the identification of *E. faecalis, E. faecium, E. hirae*, and *E. avium*, respectively. Strikingly, *E. gallinarum* was not identified by using *16S rRNA* sequencing, while the identification rate using *E. gallinarum* – LAMP and *E. gallinarum-PCR* was 100%. This indicated that *16S rRNA* sequencing was less efficient in identifying the *Enterococcus* species than LAMP assays and conventional PCR. Finally, VITEK 2 system was used for the detection of three *E. cecorum* and ten *S. aureus* strains, and the congruence of the LAMP assays and VITEK 2 was 100% for both bacteria (data not shown). The results of the nine types of LAMP assays were also confirmed by sequencing. This indicated that the LAMP assays yielded accurate results within 30 min compared with those generated by *16S rRNA* sequencing (18 to 24 h), VITEK 2 (18 to 24 h), and conventional PCR (3 to 4 h). Further, the high specificity and amplification ability of the ten types of LAMP assays allowed an easy and rapid visualization of the amplification success without the need for gel electrophoresis.

We presented the first-ever Entero-Common-LAMP assay for a simultaneous detection of eight *Enterococcus sp*. such as *E. faecalis, E. faecium, E. hirae, E. gallinarum, E. avium, E. durans, E. cecorum*, and *E. columbae*, and seven types of Entero-Specific-LAMP assays for a specific identification of seven *Enterococcus sp*. such as *E. faecalis, E. faecium, E. hirae, E. gallinarum, E. avium, E. durans*, and *E. cecorum*, using new target genes (10).

*Enterococcus sp., E. coli*, and *S. aureus* strains that cause lameness are widely found in warm-blooded animals and several other natural habitats, but are difficult to control in animals. Furthermore, *E. faecalis, E. faecium*, and *S. aureus* found in human and processed foods are major pathogens in hospital-acquired and community-acquired infection (3, 18).

In conclusion, the Entero-Common-LAMP, seven types of Entero-Specific-LAMP assays, *E. coli*-LAMP, and *S. aureus*-LAMP established herein have the potential to become a very useful tool for the prevention of disease transmission or outbreaks in the field or in resource-limited environments.

## ACKNOWLEDGMENTS

This work was supported by a research grant (B-1543084-2016-17-04) from the Animal and Plant Quarantine Agency (QIA), Ministry of Agriculture, Food and Rural Affairs, the Republic of Korea.

